# A Benchmark for Generative Modeling of High-Dimensional Enzyme Kinetic Parameters of Metabolic Networks

**DOI:** 10.64898/2026.11.25.769162

**Authors:** Stefanos Xenios, Alexandros Doulaveris, Konstantinos Mexis, Antonis Kokossis

**Affiliations:** Chemical Engineering, National Technical University of Athens, Iroon Polytechneiou 9, 15772, Athens, Greece

**Keywords:** kinetic modeling, genome-scale models, variational autoencoder, generative adversarial network, normalizing flow, benchmark, DBTL, generative models

## Abstract

Large-scale kinetic models predict how metabolism responds to genetic and environmental perturbations, but building them at scale is limited by the scarcity of experimentally determined enzyme kinetic parameters. Generative machine learning has been proposed to address this bottleneck, yet prior work has each introduced a single, bespoke architecture, leaving open which generative paradigm suits this problem and why. We introduce a benchmark suite for generative modeling of high-dimensional enzyme kinetic parameter sets: labeled datasets of kinetically feasible Escherichia coli models paired with a mechanistically grounded evaluation harness that scores generated ensembles on dynamical stability and agreement with experimentally observed phenotypes, rather than statistical realism alone. Using it, we compare three standard architectures at matched capacity, a conditional *β*-variational autoencoder, a Wasserstein GAN with gradient penalty, and a RealNVP normalizing flow, each under two conditioning schemes (a discrete high/low label and a continuous, graded key-performance-indicator target), across a difficulty ladder from one mutant phenotype to a stringent class of nine reproduced jointly. Two findings stand out. The architectures trace a consistent concentration–exploration trade-off, the VAE concentrating, the flow exploring and extrapolating, the GAN in between, but this ordering is overturned at high stringency by the conditioning scheme: at the joint class, continuous targets reach 33–117-fold enrichment over random sampling for every architecture, whereas discrete labels stay at 1.3–15-fold. Beyond the comparison, the suite is an accessible, mechanistically validated reference task for groups outside the kinetic-modeling community to benchmark their own generative algorithms.

## Introduction

Kinetic models capture the dynamics of metabolite concentrations, fluxes, and enzyme levels, and can therefore predict how a metabolic network responds to genetic and environmental perturbations in a way that stoichiometric, steady-state reconstructions cannot (Heinrich and Schuster, 1996; Cornish-Bowden, 2012; Liebermeister et al., 2010). Their construction at genome scale, however, is fundamentally limited by the scarcity of experimentally determined kinetic parameters, such as Michaelis–Menten constants and catalytic rate constants, for most organisms and conditions of interest (Liebermeister et al., 2010; Chakrabarti et al., 2013; Stanford et al., 2013). Traditional parameterization strategies, including Monte Carlo sampling within uncertainty-based frameworks such as ORACLE (Miskovic and Hatzimanikatis, 2010), can produce large ensembles of kinetically feasible models, but only a small fraction of randomly sampled parameter sets are typically both dynamically stable and physiologically plausible, and an even smaller fraction reproduce specific experimentally observed perturbation responses (Stanford et al., 2013).

Generative machine learning has recently been proposed to accelerate this process. REKINDLE (Choudhury et al., 2022) uses a generative adversarial network to learn from kinetic models that are stable and robust around a reference steady state. RENAISSANCE (Choudhury et al., 2024) instead trains a generator without labeled data, using natural evolution strategies driven by a scoring function, to reach large-scale kinetic models with dynamic properties matching experimental observations; exploring the latent space of such generators gives control over the dynamics of the generated models (Choudhury et al., 2026). Our own prior work, GENKI (Xenios et al., 2026), introduced a conditional variational autoencoder trained on phenotype-based KPIs derived from multi-omics and bioreactor data, so that generated parameter sets are enriched not merely for stability but for agreement with mutant- and condition-specific experimental observations.

Each of these methods, however, commits to a single generative architecture. What is missing is a systematic answer to a more basic question: for the specific structure of this problem, high-dimensional, tabular, non-i.i.d. kinetic parameter vectors, labeled by a mechanistic, simulation-derived performance score rather than a natural image or text distribution, which generative modeling paradigm is best suited, and along which axes do the standard architectures differ? The broader machine learning literature on generative model benchmarking (e.g., for tabular data) has developed a rich toolkit of statistical fidelity and diversity metrics (Kynkäänniemi et al., 2019; Naeem et al., 2020; Alaa et al., 2022), but these metrics evaluate distributional realism against a real dataset, not whether a generated sample corresponds to a mechanistically valid, phenotypically correct dynamical system when simulated. A recent position paper has further shown that current statistical fidelity and diversity metrics all fail some basic sanity checks (Räisä et al., 2025), reinforcing the need for a complementary, mechanistically grounded evaluation axis in any domain, like this one, where a downstream simulator can supply it.

This work makes two contributions. First, we introduce a benchmark suite, labeled datasets of kinetically feasible *E. coli* models together with the KPI-based evaluation harness developed for GENKI, that packages this mechanistic validation into a form usable by researchers outside the kinetic-modeling community. Second, we use this harness to run a controlled, matched-capacity comparison of three standard generative architectures representative of the field’s main paradigms: a conditional *β*-variational autoencoder (Higgins et al., 2017; Sohn et al., 2015), a conditional Wasserstein GAN with gradient penalty (Mirza and Osindero, 2014; Arjovsky et al., 2017; Gulrajani et al., 2017), and a conditional RealNVP normalizing flow (Dinh et al., 2017; Rezende and Mohamed, 2015).

Benchmark scenarios are organized as a difficulty ladder of increasing perturbation multiplicity within *E. coli* central carbon metabolism, from a single mutant phenotype to a maximally stringent, difficulty-calibrated joint class over nine mutant phenotypes (see *Benchmark Scenarios*).

## Methods

### Benchmark Framework Overview

The benchmark evaluates generative architectures along a four-stage pipeline (Fig. 1): (a) conditional generative models are trained on KPI-labeled kinetic parameter ensembles and used to synthesize new parameter sets targeted at a high-performance conditioning label; (b) generated parameter sets are subjected to stability-based filtering via Jacobian analysis, retaining only models that are both locally stable and physiologically fast relative to the reference doubling time; (c) surviving models are simulated under the benchmark’s perturbation protocol and scored with the KPI framework; (d) the resulting scores are summarized with the statistical, biological, and computational metrics described under *Evaluation Framework*, and compared across architectures and against the ORACLE baseline ensemble.

**Figure 1.**
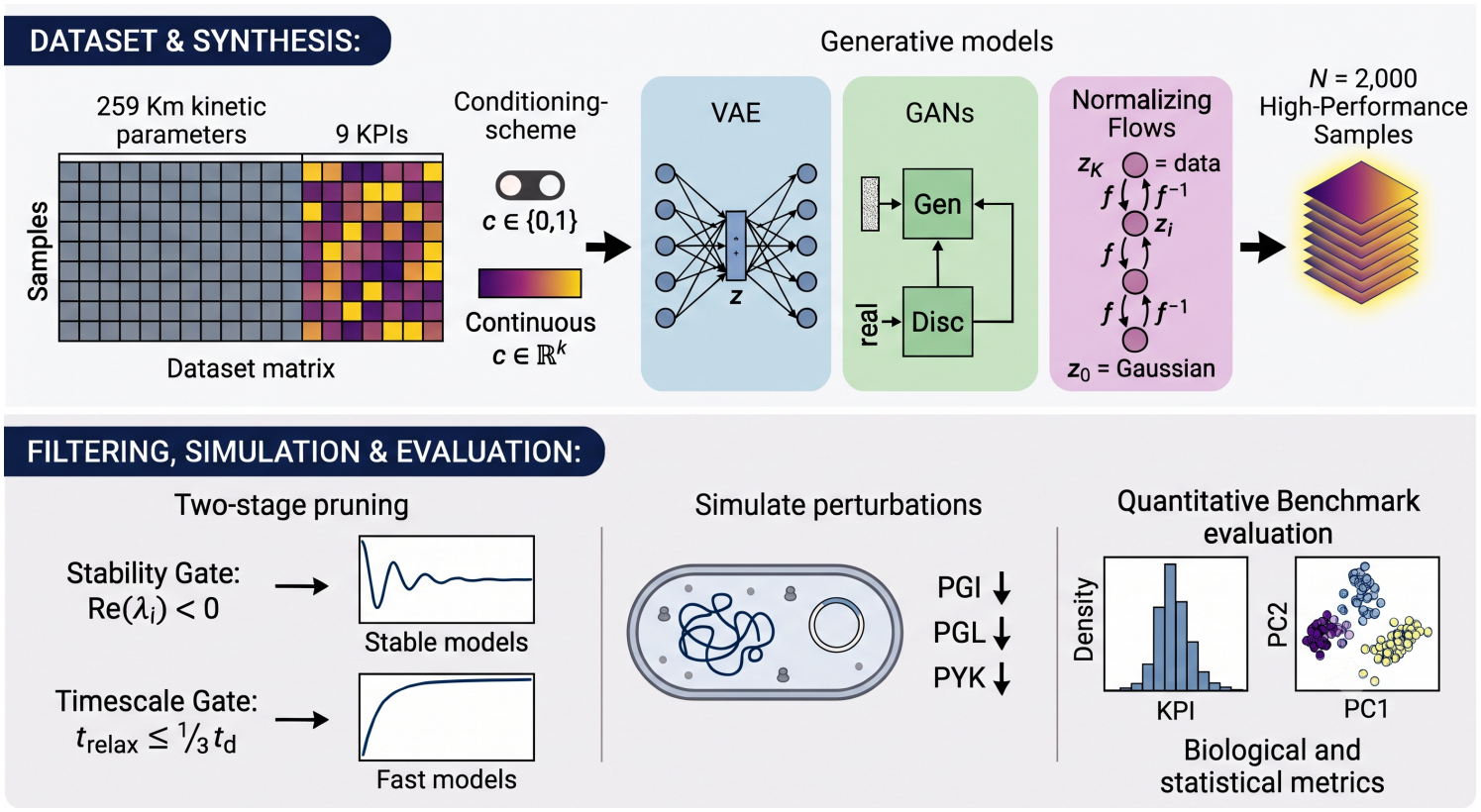
Overview of the benchmark and KPI-based evaluation framework. Top row: KPI-labeled dataset, conditioning scheme, and conditional generation of kinetic parameters with a VAE, GAN, and normalizing flow. Bottom row, left to right: Jacobian-based stability filtering; dynamic simulation and KPI scoring against experimental references; statistical and biological evaluation of the generated ensembles.

### Benchmark Dataset Construction

The benchmark dataset consists of high-dimensional enzyme kinetic parameter (*K*_*m*_) vectors and their associated KPI values for the *Escherichia coli* core metabolic network. The dataset was generated using Monte Carlo sampling within the ORACLE framework (Miskovic and Hatzimanikatis, 2010), as implemented in the skimpy software package (Weilandt et al., 2023), constrained around the reference aerobic physiology of Ishii et al. (2007). Only the Michaelis constants are modeled; for every parameter set, ORACLE or generated, the maximal velocities are back-calculated from the reference fluxes, so each *K*_*m*_ vector defines a complete model at the reference steady state. After the stability screen below, the dataset holds about 12,500 feasible models, each a 259-dimensional *K*_*m*_ vector labeled with nine KPIs, one per simulated mutant (Supplementary Tables S1–S2), log-transformed and standardized before training.

To enable supervised conditional guidance, two labelling schemes were tested. In the *discrete* scheme, per-scenario KPI values were converted into binary labels by percentile-based thresholding, computed independently per scenario and fixed before training to avoid data leakage: sets below the scenario-specific threshold were labeled high-performing (label 1), all others low-performing (label 0). In the *continuous* scheme, the graded log-KPI values themselves (standardized, one entry per perturbation) served directly as conditioning targets. *Benchmark Scenarios* specifies the thresholds per scenario, including the difficulty-calibrated per-perturbation thresholds of the stringent joint class.

### Key Performance Indicator (KPI) Framework

Model performance is quantified with the composite, ORACLE-normalized KPI (*ζ*-score) introduced in GENKI (Xenios et al., 2026), rather than a flat pooled error metric, so that no single large-magnitude flux dominates the score and each observable category (growth, uptake, secretion, intracellular, and perturbation-specific fluxes) contributes with equal structural weight regardless of how many observables it contains:

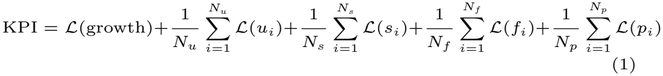

where *u*_*i*_, *s*_*i*_, *f*_*i*_, and *p*_*i*_ denote uptake, secretion, intracellular, and perturbed fluxes respectively, *N*_*u*_, *N*_*s*_, *N*_*f*_, *N*_*p*_ are the corresponding observable counts, and *ℒ* is the per-observable loss function. We use the *ζ*-score as *ℒ*, a data-driven, scale-free error function constructed from the baseline ORACLE ensemble:

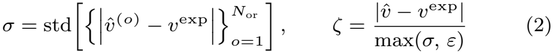

where *ε* is a small fixed floor on the denominator (set relative to the reference glucose uptake rate; see Supplementary Material), *σ* is the reaction-specific, entirely data-driven spread of ORACLE residuals for that observable, and *ζ* is interpreted as the number of ORACLE-error standard deviations by which a candidate model’s prediction departs from the experimental reference.

### Secondary diagnostic metrics: R^2^ and rank correlation

As a secondary, diagnostic complement to the KPI, we report the coefficient of determination (*R*^2^) between simulated and experimentally observed fluxes on ensemble-mean predictions. *R*^2^ is computed on log-transformed flux errors.

### Generative Architectures

Three standard conditional generative architectures were trained on the KPI-labeled datasets described above: a conditional *β*-VAE (cVAE) (Higgins et al., 2017; Sohn et al., 2015), a conditional WGAN-GP (cGAN) (Mirza and Osindero, 2014; Arjovsky et al., 2017; Gulrajani et al., 2017), and a conditional RealNVP normalizing flow(NF) (Dinh et al., 2017; Rezende and Mohamed, 2015); the suffix “-cont” denotes the continuous conditioning scheme. Each architecture is trained to model the conditional distribution of kinetic parameter sets given a performance target *c*, and subsequently to generate high-performing ensembles. A separate model is trained for every scenario and random seed; the seed fixes both network initialization and sampling. Architectures were intentionally kept close to their standard, “vanilla” form to enable an unbiased comparison and to keep the benchmark easy for other groups to reproduce or extend with new architectures. Network depth, trainable parameter count, and training hyperparameters (batch size, learning-rate schedule, epochs) were matched as closely as possible across the three models (Supplementary Table S3). Architecture-specific exploration/fidelity controls, such as VAE sampling temperature, GAN truncation, or normalizing-flow temperature, were deliberately *not* swept or tuned per architecture; each model was run at its standard default operating point (in particular, the VAE and flow sampling temperature was fixed at 1), so that differences observed between architectures reflect each paradigm’s inherent inductive bias rather than the degree of tuning invested in it. A full description of each architecture, together with its complete mathematical formulation, the two conditioning schemes (discrete and continuous), and the training configuration, is given in the Supplementary Material; continuous targets use the same thresholds as the discrete label.

### Model Screening: Stability-Based Filtering

Before KPI evaluation, every generated kinetic parameter set was subjected to the same two-stage screening procedure used to construct the baseline ORACLE ensemble.

#### Dynamic stability

The Jacobian matrix of the ODE system is evaluated at the reference steady state; only parameter sets whose Jacobian eigenvalues all have strictly negative real parts are retained.

#### Physiological timescale

The characteristic relaxation time, estimated from the dominant Jacobian eigenvalue, must be at least three times faster than the organism’s doubling time *t*_*d*_ = ln 2*/µ*, excluding models that are mathematically stable but unrealistically slow.

### Dynamic Simulation and Phenotype Evaluation

Models surviving stability-based filtering were simulated under the benchmark’s perturbation protocol (see *Benchmark Scenarios*) and scored with the KPI framework above, using the same experimental references used to construct the baseline ensemble. The same simulation and scoring procedure is applied identically to the baseline ensemble and all three generative architectures, ensuring a fair, consistent comparison. Full specifics of the dynamic simulation protocol are provided in the Supplementary Material.

### Evaluation Framework

Evaluation metrics are organized into two families—biological/phenotypic and statistical fidelity/diversity—so that the Results can report one leaderboard table per scenario with one column block per family, rather than a flat, unstructured metric list. The primary metric is the *enrichment factor*, the fraction of screened models in the targeted high-performance class divided by the same fraction for ORACLE; all other metrics are diagnostics of *how* a method attains it.

#### Biological / phenotypic metrics

We report per ensemble the mean KPI (Eq. 1); the percentage of models satisfying the conditional KPI threshold used during training (whether the generator reproduces the targeted high-performance regime); the percentage outperforming the best original (ORACLE) sample; the enrichment factor for models within the desired class; the stability pass rate; and the *R*^2^ between simulated and experimental fluxes in log space. All percentages other than the pass rate are computed, for ORACLE and every generator alike, over the models that passed the screen.

#### Statistical fidelity and diversity metrics

As a complement to the biological metrics above (and because statistical realism alone is not a sufficient proxy for biological validity), we characterize each generated ensemble in parameter space with three sample-based quantities computed in a shared, PCA-reduced feature space: *precision*, the fraction of generated parameter sets inside the reference manifold (fidelity); *recall*, the fraction of reference sets inside the generated manifold (coverage/diversity) (Kynkäänniemi et al., 2019); and the *extrapolation distance* 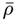, a normalized nearest-neighbour distance measuring how far generatedsets venture beyond the reference (*ρ* ≈ 1 interpolation, *ρ* ≫ 1 genuine extrapolation, *ρ* ≈ 0 near-duplication/memorization). All statistical metrics are reported as the mean *±* standard deviation across random seeds. Full definitions, the feature-space construction, and the exact formulae are given in the Supplementary Material.

### Benchmark Scenarios

Each perturbation reproduces one single-gene deletion mutant of Ishii et al. (2007), modeled as the proteomics-measured reduction in the corresponding enzyme’s abundance (Supplementary Table S2). Mutants are simulated independently, so a multi-perturbation scenario asks one parameter set to reproduce several *single*-mutant phenotypes at once, not a combinatorial multiple mutant. Per-perturbation thresholds are calibrated to individual difficulty so that the joint class stays populated but grows rarer up the ladder.

The ladder comprises three rungs: *Scenario I*, the PGI mutant alone (baseline); *Scenario II*, the PGI and PGL mutants jointly, with independent per-perturbation thresholds (lowest 10% for PGI, 30% for PGL); and *Scenario III*, a maximally stringent joint class over all nine mutants at difficulty-calibrated thresholds.

## Results

### Scenario I: PGI Mutant

For each of the six variants (three architectures *×* two conditioning schemes), 2,000 samples were generated under the high-performanceregime, filtered for stability, simulated, and scored over 4 seeds (3 for cGAN-cont); full metrics are in Supplementary Table S4.

Every variant improved on the ORACLE baseline by a wide margin (mean KPI 12.05 → 7.4–9.7; in-threshold fraction 10% → 33–88%), but the families differ in *how* they enrich, tracing aclear concentration–exploration axis (Fig. 2): the cVAE concentrates (lowest mean KPI 7.45, highest in-threshold 87.5%, 100% stability, but recall 0.00 and 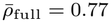), the normalizing flow explores (recall 0.37, genuine extrapolation 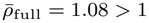, and the largest fractionbeyond the best ORACLE model, at the lowest stability ∼ 74%), and the cGAN lies between. Crucially, continuous conditioning confers essentially no advantage at this easy single-mutant scenario—each continuous variant tracks its discrete counterpart on every metric—because the high-performance class is common (∼ 10% of ORACLE). Scenario I is therefore the informative baseline of the difficulty ladderrather than a differentiator between conditioning schemes; the six variants also converge on the same underlying high-performance manifold, with a shared sign structure of log(*K*_*m*_) shifts across architectures (Supplementary Fig. S1).

**Figure 2.**
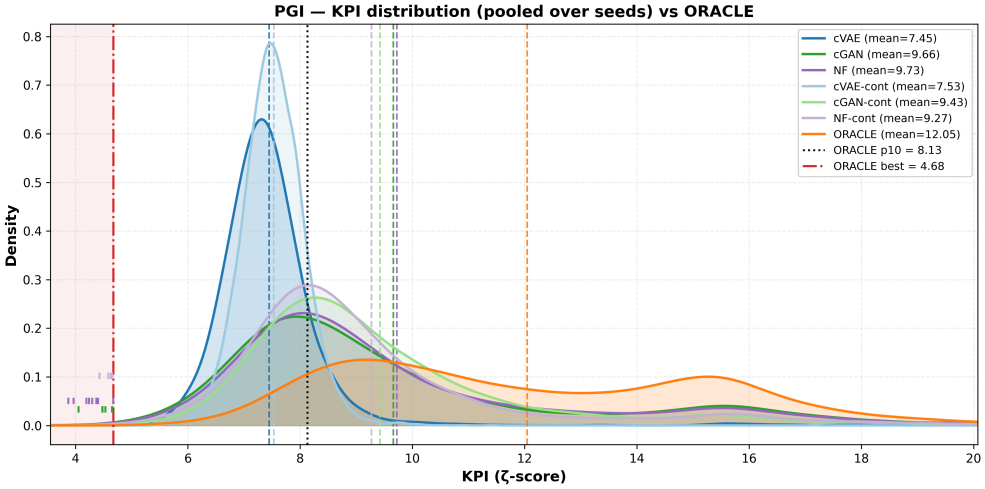
Scenario I (PGI), pooled over seeds. KPI (*ζ*-score) densities (dotted: ORACLE p10; dash-dot: best ORACLE).

### Scenario II: PGI and PGL Mutants Jointly

#### Partition design

The second rung of the ladder requires a model to satisfy *two* perturbation constraints simultaneously. The two perturbations are partitioned at different stringencies: PGI, the harder and more mechanistically constrained perturbation, is held to the stronger partition (its lowest 10% of KPI values define the high-performance class), while the more permissive PGL is partitioned at its lowest 30%. A model is counted as jointly high-performing only if it falls below *both* thresholds. This asymmetric partitioning keeps the joint class populated yet rare: only 3.1% of the baseline ORACLE ensemble satisfies both constraints at once (versus 10% and 30% for PGI andPGL alone; 0.1*×*0.3 = 3%, i.e. the two KPIs are nearly independent), the base rate against which enrichment is measured (Table 1).

**Table 1.**
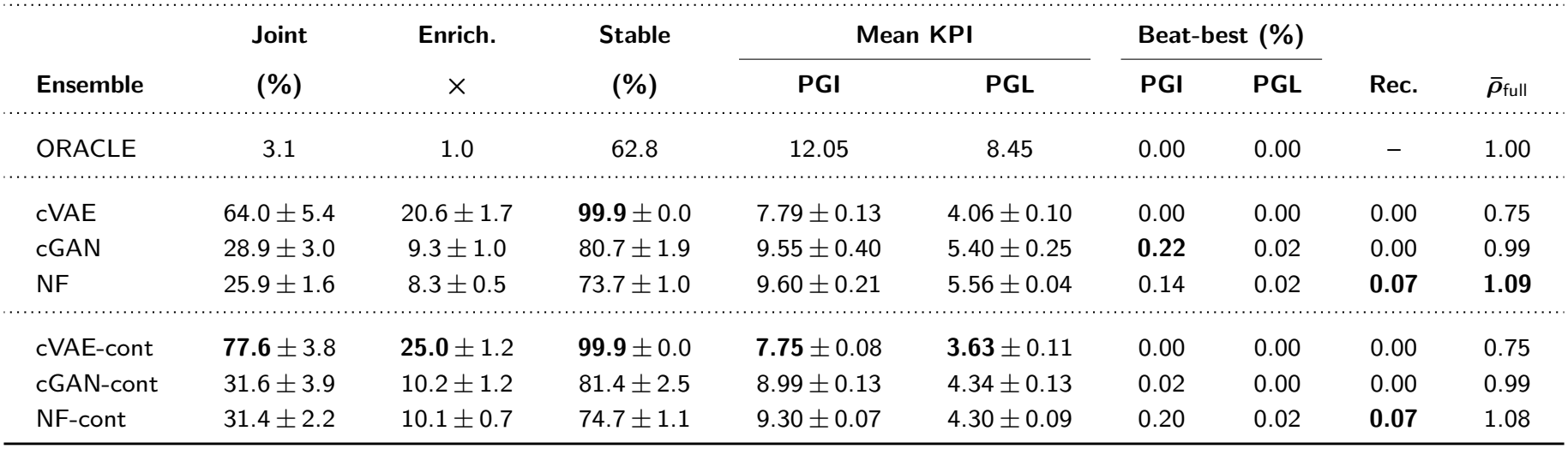
Scenario II (PGI+PGL) leaderboard; mean *±* s.d. over seeds (*n* = 3). Joint %: fraction below both partitions; Enrich.: over ORACLE base rate (3.1%); Stable: pass rate; mean KPI and Beat-best per perturbation; Rec.: coverage (Kynkäänniemi et al., 2019); 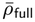: mean NN distance to ORACLE. Best per column (excl. ORACLE) in bold.

| **Ensemble** | **Joint** | **Enrich.** | **Stable** | **Mean KPI** |  | **Beat-best (%)** |  |  |  |
|---|---|---|---|---|---|---|---|---|---|
|  | **(%)** | × | **(%)** | **PGI** | **PGL** | **PGI** | **PGL** | **Rec.** | ***ρ̄***<sub>full</sub> |
| ORACLE | 3.1 | 1.0 | 62.8 | 12.05 | 8.45 | 0.00 | 0.00 | – | 1.00 |
| cVAE | 64.0 ± 5.4 | 20.6 ± 1.7 | **99.9** ± 0.0 | 7.79 ± 0.13 | 4.06 ± 0.10 | 0.00 | 0.00 | 0.00 | 0.75 |
| cGAN | 28.9 ± 3.0 | 9.3 ± 1.0 | 80.7 ± 1.9 | 9.55 ± 0.40 | 5.40 ± 0.25 | **0.22** | 0.02 | 0.00 | 0.99 |
| NF | 25.9 ± 1.6 | 8.3 ± 0.5 | 73.7 ± 1.0 | 9.60 ± 0.21 | 5.56 ± 0.04 | 0.14 | 0.02 | **0.07** | **1.09** |
| cVAE-cont | **77.6** ± 3.8 | **25.0** ± 1.2 | **99.9** ± 0.0 | **7.75** ± 0.08 | **3.63** ± 0.11 | 0.00 | 0.00 | 0.00 | 0.75 |
| cGAN-cont | 31.6 ± 3.9 | 10.2 ± 1.2 | 81.4 ± 2.5 | 8.99 ± 0.13 | 4.34 ± 0.13 | 0.02 | 0.00 | 0.00 | 0.99 |
| NF-cont | 31.4 ± 2.2 | 10.1 ± 0.7 | 74.7 ± 1.1 | 9.30 ± 0.07 | 4.30 ± 0.09 | 0.20 | 0.02 | **0.07** | 1.08 |

Every variant concentrated strongly into this rarer joint class, enriching over the ORACLE base rate by 8–25*×* (Table 1, Fig. 3). The same concentration–exploration ordering from Scenario I holds: thecVAE variants drive both per-perturbation mean KPIs lowest (PGI ≈ 7.8, PGL ≈ 3.6–4.1 versus ORACLE 12.05 and 8.45; Supplementary Fig. S2) and reach the highest joint yield (cVAE 64.0%, 20.6*×*), while the flows retain coverage (recall 0.07) and extrapolation 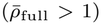 at a lower joint yield. The joint-KPI scatter (Fig. 3) makes the mechanism visual: the cVAE variants collapse almost entirely into the good-fit box, whereas the cGAN and NF clouds retain a broad tail along the harder PGI axis, and ORACLE fills the plane.

**Figure 3.**
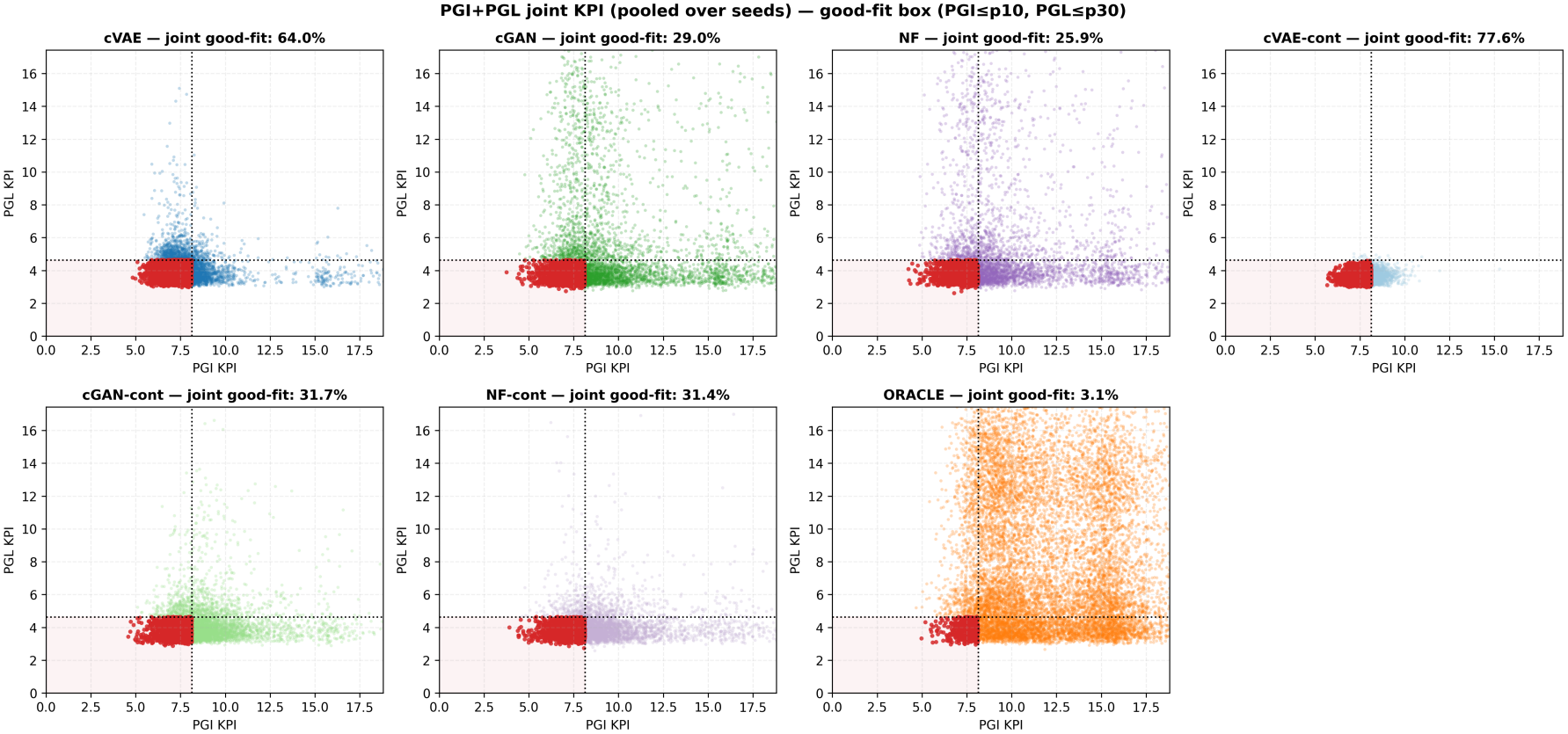
Scenario II (PGI+PGL), pooled over seeds. Joint scatter (PGL vs. PGI KPI); shaded box = joint good-fit (PGI≤p10, PGL≤p30). cVAE variants collapse into the box (up to 77.6%), cGAN/NF keep a tail on the harder PGI axis, ORACLE fills the plane (3.1%).

#### Reaching beyond ORACLE

Crucially, the two families differ in whether they produce models *better* than anything ORACLE sampled. On the harder PGI axis, only the exploratory cGAN and NF variants place any models beyond the single best ORACLE model (cGAN 0.22%, NF 0.14%, NF-cont 0.20%), while both cVAE variants produce none; on the easier PGL axis essentially no variant beats the ORACLE best. In other words, the concentrating cVAE wins on *bulk* enrichment but never sets a new record, whereas the explorers are the only methods that occasionally exceed the reference frontier. Continuous conditioning, which was essentially inert in Scenario I, also begins to pay off here (every familyimproves its joint enrichment: cVAE 20.6 → 25.0*×*, cGAN 9.3 → 10.2*×*, NF 8.3 → 10.1*×*), the first sign of the lever that becomes dominant at the capstone.

### Scenario III: Maximally Stringent Joint Class

The capstone of the benchmark asks every variant to generate models that jointly reproduce *all nine* available mutant phenotypes at once (PGI, PGL, RPI, TALA, PYK, G6PDH2r, PPS, TKT2, GND), each under a per-perturbation threshold calibrated to its individual difficulty (PGI 25, PGL 40, TALA 60, G6PDH2r 25, RPI 60, PYK 60,

PPS 40, TKT2 60, GND 40; percentile of ORACLE KPI). This is an extreme, deliberately rare target: only 0.13% of the baseline ORACLE ensemble clears all nine constraints simultaneously (16 of 12,429 training models), a base rate roughly two orders of magnitude smaller than in Scenario II.

#### Continuous conditioning is the deciding factor

At this stringency the discrete variants essentially collapse: discrete cVAE, cGAN, and NF reach the joint class at only 0.17–1.95% (1.3–15*×* enrichment), i.e. the concentrating cVAE that dominated Scenarios I–II now performs no better than random ORACLE sampling (1.3*×*). With the continuous target drawn from the same positive class, *every* architecture instead reaches 33–117*×* enrichment (Table 2, Fig. 4): cVAE-cont 15.0% (117*×*), NF-cont 5.6% (43*×*),and cGAN-cont 4.2% (33*×*), i.e. 90, 24, and 3*×* their discretecounterparts. On the softer “≥ 7 of 9” criterion the continuous variants place 40–85% of models versus 8–26% for their discretecounterparts. This is the central result of the benchmark: the graded performance target, which was inert when the good class was common (Scenario I) and mildly helpful when it was moderately rare (Scenario II), becomes decisive once the joint class is genuinely rare, and it does so across all three architecture families rather than a single one. The gap between average-case and joint performance underscores why this is the hard bar: discrete NF is the strongest discrete method on mean KPI averaged over the nine perturbations (9.81, Table 2), yet clears all nine thresholds *simultaneously* in only 1.95% of models—being close on each perturbation on average is not the same as satisfying every one at once, which is precisely the capability continuous conditioning unlocks. A graded target can also be pushed beyond the positive class: at the 5^th^ KPI percentile, enrichment rose to 95–416*×* (Supplementary Table S5, Fig. S5).

**Table 2.**
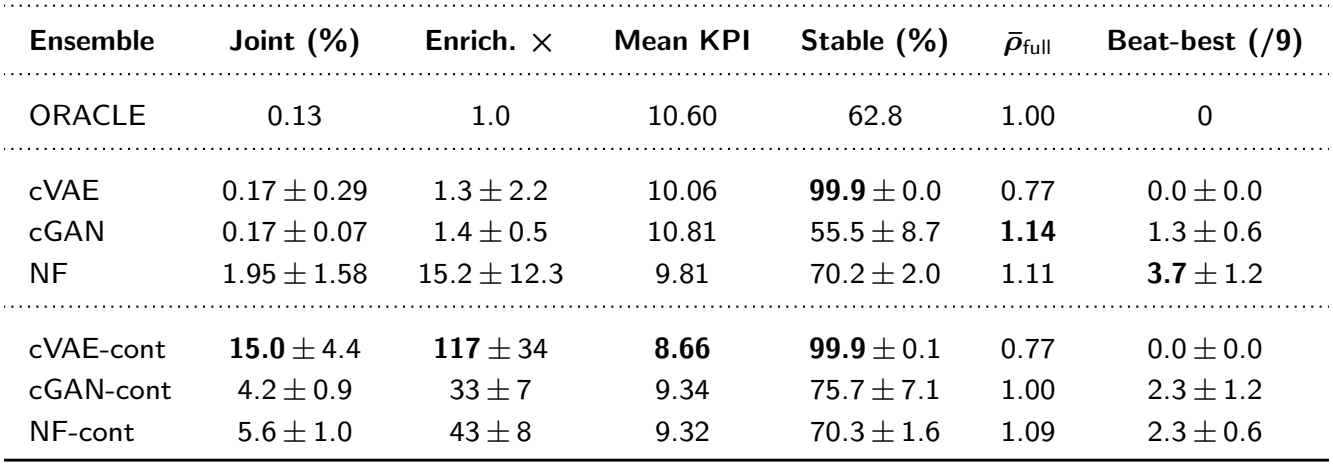
Scenario III (9-mutant joint class) leaderboard; mean *±* s.d. over seeds (*n* = 3). Joint %: fraction satisfying all nine thresholds; Enrich.: over ORACLE base rate (0.13%); mean KPI averaged over the nine perturbations (lower better; 3 non-physical KPIs *>* 10^3^ excluded); Stable: pass rate; 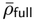: mean NN distance to ORACLE; Beat-best: perturbations (of nine) with a model beating the best ORACLE model. Best per column (excl. ORACLE) in bold.

| **Ensemble** | **Joint (%)** | **Enrich. ×** | **Mean KPI** | **Stable (%)** | $\bar{\rho}_{\text{full}}$ | **Beat-best (/9)** |
|---|---|---|---|---|---|---|
| ORACLE | 0.13 | 1.0 | 10.60 | 62.8 | 1.00 | 0 |
| cVAE | 0.17 ± 0.29 | 1.3 ± 2.2 | 10.06 | **99.9** ± 0.0 | 0.77 | 0.0 ± 0.0 |
| cGAN | 0.17 ± 0.07 | 1.4 ± 0.5 | 10.81 | 55.5 ± 8.7 | **1.14** | 1.3 ± 0.6 |
| NF | 1.95 ± 1.58 | 15.2 ± 12.3 | 9.81 | 70.2 ± 2.0 | 1.11 | **3.7** ± 1.2 |
| cVAE-cont | **15.0** ± 4.4 | **117** ± 34 | **8.66** | **99.9** ± 0.1 | 0.77 | 0.0 ± 0.0 |
| cGAN-cont | 4.2 ± 0.9 | 33 ± 7 | 9.34 | 75.7 ± 7.1 | 1.00 | 2.3 ± 1.2 |
| NF-cont | 5.6 ± 1.0 | 43 ± 8 | 9.32 | 70.3 ± 1.6 | 1.09 | 2.3 ± 0.6 |

**Figure 4.**
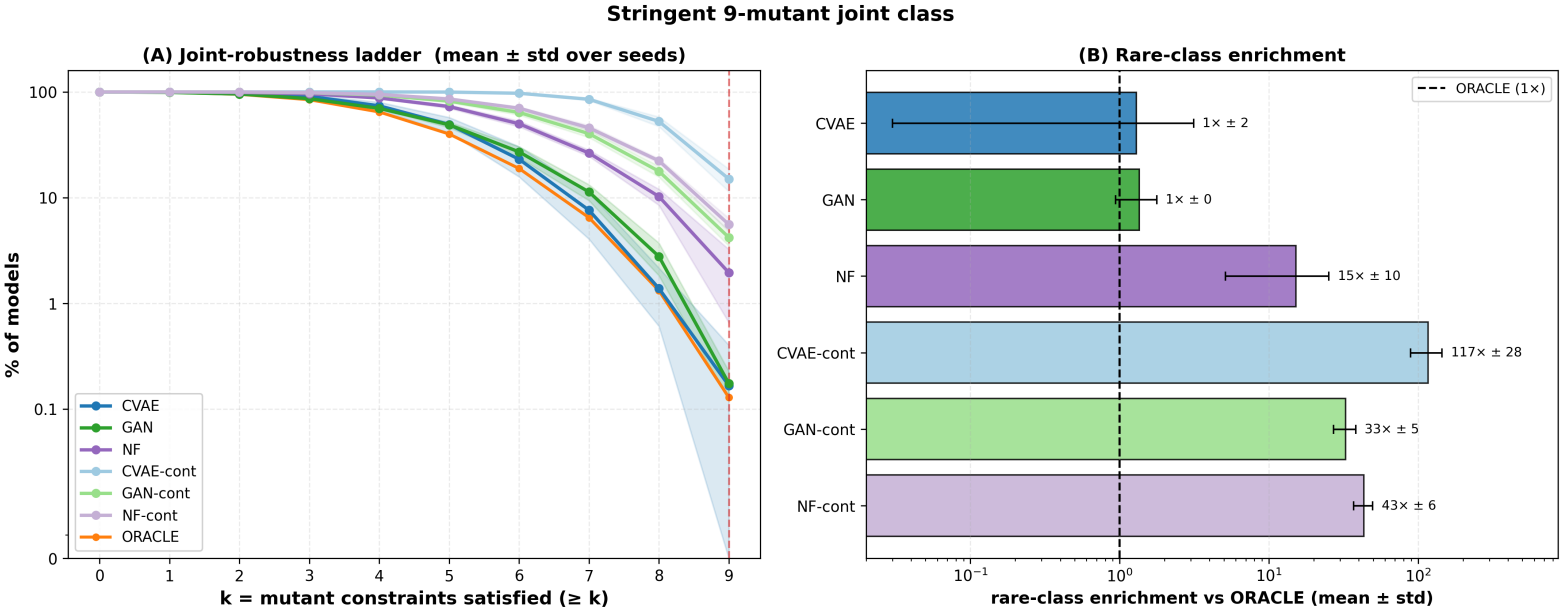
Scenario III (9-mutant joint class), pooled over seeds. Joint-robustness ladder (% of models satisfying ≥ *k* of 9 thresholds) and rare-class enrichment vs. ORACLE; continuous variants use positive-class targets and reach 33–117*×* (discrete: 1.3–15*×*). Per-mutant densities in Supplementary Fig. S3.

## Discussion

The three rungs of the difficulty ladder jointly expose the distinct inductive bias of each generative architecture and the decisive role of the conditioning scheme. Across every conditioning scenario, thearchitectures trace a consistent concentration–exploration axis: the conditional VAE concentrates almost perfectly within the region on which it is conditioned, attaining the lowest mean KPI and the highest in-threshold yields but essentially no coverage beyond the reference; the normalizing flow extrapolates furthest 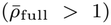, which is why it most frequently produces models that surpass the best member of the initial ORACLE ensemble; and the GAN occupies the intermediate position, capable of setting new best models while still concentrating efficiently. This ordering is stable and holds under both discrete and continuous conditioning.

The conditioning signal governs whether the generators can reproduce stringent, and potentially unseen, performance classes. When the target class is common or only moderately rare, continuous conditioning confers little advantage, because it asks the generator to hit the graded target at that exact percentile, whereas the discrete label additionally carries the information of all models lying far below the threshold. The balance inverts once the class becomes genuinely stringent: the discrete label collapses to a single, near-empty category that is too underrepresented for any architecture to learn the parameter patterns that generated it, while the continuous scheme retains the full, graded performance information of every experiment rather than a flat binary tag and can interpolate toward the joint optimum. Two properties drive the collapse: the positive class holds only 16 training examples, and the discrete label compresses nine per-mutant outcomes into one bit, whereas the continuous target carries nine graded KPIs; a per-mutant multi-label scheme would separate these effects. Because this reversal appears identically across the variational, adversarial, and flow families, we conclude that it is a property of the conditioning signal itself rather than of any one architecture.

We paired the mechanistic KPI with statistical fidelity and diversity metrics precisely because the latter dominate tabular generative-model evaluation yet need not track mechanistic validity (Kynkäänniemi et al., 2019; Räisä et al., 2025). Although the generators may appear merely to copy the original dataset, mechanistic KPI enrichment spans more than an order of magnitude across those same variants. A generator can thus be statistically almost indistinguishable from the reference while being up to two orders of magnitude better at producing simulator-valid, phenotype-matching models, which supports treating the KPI as the primary axis of comparison and the statistical metrics as a complementary diagnostic.

## Conclusion

We introduced a benchmark suite for generative modeling of high-dimensional enzyme kinetic parameters—kinetically feasible *E. coli* ensembles paired with a mechanistically grounded, GENKI-derived KPI harness—and used it to compare a conditional VAE, GAN, and normalizing flow under discrete and continuous conditioning across a difficulty ladder from one to nine jointly reproduced mutant phenotypes. The architectures trace a stable concentration– exploration axis, but this ordering is overturned at high stringency by the conditioning scheme: at the joint class, continuous, gradedKPI targets reach 33–117*×* enrichment over ORACLE for every architecture where discrete labels stay at 1.3–15*×*—a cross-family effect only a shared-harness comparison could establish. Statistical fidelity, meanwhile, does not predict mechanistic performance. Beyond these findings, the suite is intended as a reusable, mechanistically validated reference task for the broader community to benchmark new generative algorithms under identical, reproducible conditions.

## Acknowledgements

This project has received funding from the European Union’s Horizon 2020 research and innovation programme BIOINDUSTRY 4.0 (grant agreement 101094287).

## Data availability

Training data and model files are included in the code repository; generated parameter sets, trained networks and simulation outputs are on Zenodo (https://doi.org/10.5281/zenodo.24532943).

## Code availability

KinGenBench is available at https://github.com/IPSEN-NTUA/KinGenBench (MIT license).

## Declaration of generative AI and AI-assisted technologies in the writing process

During the preparation of this work, the authors used AI-assisted tools to help draft and restructure parts of the manuscript. The authors reviewed and edited the output as necessary and take full responsibility for the content of the publication.

