## Supplementary material for "A Benchmark for Generative Modeling of High-Dimensional Enzyme Kinetic Parameters of Metabolic Networks"

*Bioinformatics*, YEAR, pp. 1–6
DOI added during production
Published: Date added during production
**Supplementary Material**

### Supplementary Material
### A Benchmark for Generative Modeling of High-Dimensional Enzyme Kinetic Parameters of Metabolic Networks

Stefanos Xenios,[1,*] Alexandros Doulaveris,[1] Konstantinos Mexis[1] and Antonis Kokossis[1]

[1]Chemical Engineering, National Technical University of Athens, Iroon Polytechneiou 9, 15772, Athens, Greece

#### Abstract

**This document provides supplementary material for the main manuscript. It covers, in order: the construction, filtering, and preprocessing of the benchmark dataset together with the in-silico perturbation experiments that define its labels (Section S1); conceptual background and the full configuration of the three conditional generative architectures and the two conditioning schemes (Section S2); the dynamic-simulation protocol used to score generated models (Section S3); the complete definitions of the statistical fidelity and diversity metrics (Section S4); and the per-scenario leaderboard and additional figures (Sections S5–S6).**

#### S1. Benchmark dataset and perturbation scenarios

##### S1.1. Dataset construction, filtering, and preprocessing

The *E. coli* benchmark dataset was generated by Monte Carlo sampling within the ORACLE framework (Miskovic and Hatzimanikatis, 2010), as implemented in the `skimpy` package (Weilandt et al., 2023), around the reference aerobic steady state of Ishii et al. (2007). Each sampled parameterization is a 259-dimensional vector of Michaelis–Menten constants ($K_m$) for the *E. coli* core network (Table S1). The $K_m$ values are the only modeled features: for every parameter set, whether sampled by ORACLE or produced by a generator, the forward maximal velocities are back-calculated in `skimpy` from the reference steady-state fluxes and concentrations, so that each $K_m$ vector defines a complete kinetic model consistent with the reference steady state. The equilibrium constants are fixed by thermodynamics and shared by all models.

Every sampled model was passed through the same two-stage screen used to build the baseline ORACLE ensemble (detailed in Section S3): a model was retained only if (i) all eigenvalues of the Jacobian of the ODE system, evaluated at the reference steady state, have strictly negative real parts (*dynamic stability*), and (ii) its characteristic relaxation time is at least three times faster than the cellular doubling time $t_d = \ln 2/\mu$ (*physiological timescale*), with $\mu = 0.2\,\mathrm{h}^{-1}$ for *E. coli* ($t_d \approx 3.47\,\mathrm{h}$). Applying this screen yields the 12,524 kinetically feasible models that constitute the benchmark training set. Because a model enters a scenario only if every perturbation simulation that scenario requires completed, the per-scenario training sets are slightly smaller: 12,503 models in Scenario I, 12,475 in Scenario II, and 12,429 in Scenario III. Under the discrete scheme the positive (jointly high-performing) class then holds 1,251 (10.0%), 388 (3.1%), and 16 (0.13%) models, respectively. Prior to training, the raw $K_m$ values were log-transformed and then standardized to zero mean and unit variance feature-wise (z-score scaling), using statistics fixed on the training set to avoid leakage.

**Table S1** *E. coli* core network and parameter summary. $K_m$: Michaelis–Menten constants (the modeled features); $v_{max}$: maximal reaction velocities; $K_{eq}$: equilibrium constants.

| Organism | Rxn | Met | $K_m$ | $v_{max}$ | $K_{eq}$ | Total |
|---|---|---|---|---|---|---|
| *E. coli* | 95 | 72 | 259 | 95 | 70 | 424 |

Each retained model is labeled with 9 KPI values, one per single-perturbation scenario available in the current release (PGI, PGL, PYK, RPI, TALA, G6PDH2r, TKT2, GND, PPS), giving a final dataset of 268 variables (259 kinetic-parameter features plus 9 KPI columns). The composite, ORACLE-normalized KPI ($\zeta$-score) is defined in the main text; the perturbation experiments that produce each KPI are described next.

##### S1.2. In-silico perturbation experiments

Mutant phenotypes were simulated by introducing enzyme-specific perturbations into each kinetic model. A perturbation is applied as a multiplicative change to the abundance parameter of the target enzyme relative to the reference state; the magnitudes are derived from proteomics data and represent the measured abundance change of the affected enzyme in the corresponding mutant strain relative to

© The Author(s) YEAR. Copyright and licence statements to be updated by the publisher during production.

**Table S2** Enzyme abundance perturbation factors for the nine benchmark perturbations. Each value is the multiplicative change applied to the target enzyme's abundance parameter relative to the reference state, derived from proteomics measurements of the corresponding mutant strain; a factor below one denotes downregulation. For G6PDH2r no direct measurement was available and the minimum detectable abundance level of the proteomics platform was used instead.

| Perturbation | PGI | PGL | RPI | TALA | PYK | G6PDH2r | PPS | GND | TKT2 |
|---|---|---|---|---|---|---|---|---|---|
| **Abundance factor** | 0.06 | 0.10 | 0.50 | 0.33 | 0.34 | 0.013 | 0.014 | 0.001 | 0.23 |

**Table S3** Architecture and training configuration of the three conditional generative models. All models operate on the same 259-dimensional, log-transformed and standardized $K_m$ vector and are conditioned on a per-scenario performance target $c$, which is either a binary high-/low-performance label ($c \in \{0, 1\}$) or a continuous, standardized log-KPI vector ($c \in \mathbb{R}^k$, one entry per perturbation). Trainable-parameter counts are derived directly from the layer definitions. Further settings: cVAE, Gaussian encoder, mean-squared-error reconstruction loss, and KL weight $\beta$ annealed linearly from 0.1 to 1.0 over the first 80 epochs; cGAN, gradient-penalty coefficient $\lambda = 10$ and five critic updates per generator update; NF, gradient-norm clipping at 5. In the discrete scheme the binary label enters each network through a learned embedding (32-dimensional for the cGAN); in the continuous scheme the standardized KPI vector is concatenated to the network input. A separate model is trained for every scenario and seed; the seed fixes initialization and sampling.

| Model | Component MLPs | Hidden widths | Latent dim | Trainable | Optimizer (lr) | Epochs / batch |
|---|---|---|---|---|---|---|
| cVAE | encoder / decoder | 256–128–128 / 128–128–256 | 80 | 275,491 | Adam ($5 \times 10^{-4}$) | 120 / 100 |
| cGAN | generator / critic | 128–256 / 256–128 | 80 | 221,956 | Adam ($5 \times 10^{-4}$) | 120 / 100 |
| NF | 6 coupling networks | ResidualNet, 64 wide, 2 blocks | 259 (bijective) | 264,190 | AdamW ($1 \times 10^{-3}$) | 200 / 512 |

the parent strain (Ishii et al., 2007). A factor below one thus encodes a downregulation: e.g. a factor of 0.06 for PGI scales that enzyme's abundance to 6% of its reference level.

The nine per-perturbation abundance factors that define the benchmark scenarios are summarized in Table S2. A factor below one denotes a downregulation; for G6PDH2r, no direct proteomics measurement was available for the corresponding gene in the mutant strain, so the minimum detectable abundance level of the proteomics platform was used instead.

For every sampled and screened kinetic parameterization, the model is initialized at the reference steady-state concentrations, the selected enzyme perturbation is applied, and the system is integrated to its post-perturbation state (Section S3). Each perturbation is simulated independently, so a single ensemble can be scored across the single-, dual-, and nine-perturbation scenarios of the difficulty ladder without re-simulation from scratch. Fluxes read out from the post-perturbation state are compared against the experimentally observed mutant phenotypes through the KPI framework of the main text.

#### S2. Generative architectures

The benchmark compares three conditional generative models, one drawn from each of the dominant paradigms in modern generative modeling: a latent-variable model trained on a variational bound (the $\beta$-VAE), an implicit model trained adversarially (the WGAN-GP), and an exact-likelihood model built from invertible maps (the RealNVP normalizing flow). All three learn the conditional distribution $p(\mathbf{x} \mid c)$ of kinetic parameter vectors $\mathbf{x}$ given a performance target $c$, but they differ in how that distribution is represented and optimized, and these differences give each a characteristic inductive bias that the benchmark is designed to expose. Each model operates on the same 259-dimensional, log-transformed and standardized $K_m$ vector; network depth, trainable-parameter count, and training schedule were matched as closely as the architectures allow (Table S3) so that observed differences reflect each paradigm's inductive bias rather than tuning effort. Below, each architecture is introduced conceptually before its formal training objective.

##### Conditional $\beta$-VAE.

The variational autoencoder (Kingma and Welling, 2014) is a latent-variable model: it posits a low-dimensional latent code $\mathbf{z}$ with a simple prior and learns a decoder $p_\theta(\mathbf{x} \mid \mathbf{z}, c)$ that maps codes to data, together with an amortized encoder $q_\phi(\mathbf{z} \mid \mathbf{x}, c)$ that approximates the intractable posterior. Because exact likelihood is unavailable, training maximizes the evidence lower bound (ELBO), which balances a reconstruction term against a Kullback–Leibler term pulling the approximate posterior toward the prior. The $\beta$-VAE (Higgins et al., 2017) reweights that KL term by a factor $\beta$: larger $\beta$ pushes the aggregate posterior closer to the prior, yielding a smoother, more regularized latent space at some cost in reconstruction fidelity. Two properties give the VAE its inductive bias for this problem. First, sampling is anchored to a unit-Gaussian prior and passed through a decoder trained to minimize squared reconstruction error, which concentrates generated samples near the high-density core of the target conditional and rarely far into its tails. Second, conditioning both encoder and decoder on $c$ (the conditional VAE, Sohn et al., 2015) lets the model reuse a shared latent geometry across performance levels. The net effect is a generator that *concentrates*: it reproduces the targeted high-performance regime with high yield but with limited coverage or extrapolation beyond the reference. Formally, encoder and decoder are trained jointly by maximizing the $\beta$-weighted ELBO

$$\mathcal{L}_{\beta\text{-VAE}}(\phi, \theta) = \mathbb{E}_{q_\phi(\mathbf{z}|\mathbf{x},c)}\big[\log p_\theta(\mathbf{x} \mid \mathbf{z}, c)\big] - \beta\,\mathrm{KL}\big(q_\phi(\mathbf{z} \mid \mathbf{x}, c)\,\|\,p(\mathbf{z})\big), \tag{S1}$$

with $q_\phi(\mathbf{z} \mid \mathbf{x}, c) = \mathcal{N}\big(\boldsymbol{\mu}_\phi(\mathbf{x}, c), \operatorname{diag}\boldsymbol{\sigma}^2_\phi(\mathbf{x}, c)\big)$, prior $p(\mathbf{z}) = \mathcal{N}(\mathbf{0}, \mathbf{I})$, and the expectation estimated via the reparameterization trick $\mathbf{z} = \boldsymbol{\mu}_\phi + \boldsymbol{\sigma}_\phi \odot \boldsymbol{\epsilon}$, $\boldsymbol{\epsilon} \sim \mathcal{N}(\mathbf{0}, \mathbf{I})$.

##### Conditional WGAN-GP.

The generative adversarial network (Goodfellow et al., 2014) takes an implicit, likelihood-free approach: a generator $G_\theta$ transforms noise into samples, while a discriminator is trained to separate generated from real data, and the generator is trained to fool it, a minimax game whose equilibrium matches the data distribution. Because there is no explicit density and no reconstruction term tying samples to the data core, GANs can place mass along the boundary of the data manifold, but the original formulation is prone to unstable training and mode collapse. The Wasserstein GAN (Arjovsky et al., 2017) replaces the discriminator with a *critic* that estimates the Earth-Mover (Wasserstein-1) distance between the real and generated distributions, giving smoother, more informative gradients; this requires the critic to be 1-Lipschitz, which we enforce with a gradient penalty (WGAN-GP, Gulrajani et al., 2017) rather than weight clipping. Conditioning both networks on $c$ (Mirza and Osindero, 2014) yields the conditional WGAN-GP used here. Its inductive bias sits *between* the VAE and the flow: with no explicit anchor to the high-density core it can and does propose models beyond the best reference sample, yet it still concentrates efficiently into the targeted class. Generator and critic are trained by optimizing

$$\min_{\theta}\max_{\psi}\ \mathbb{E}_{\mathbf{x}\sim p_{\mathrm{data}}}\big[D_\psi(\mathbf{x},c)\big]-\mathbb{E}_{\mathbf{z}\sim p(\mathbf{z})}\big[D_\psi(G_\theta(\mathbf{z},c),c)\big] \\ -\lambda\,\mathbb{E}_{\hat{\mathbf{x}}}\Big[\big(\|\nabla_{\hat{\mathbf{x}}}D_\psi(\hat{\mathbf{x}},c)\|_2-1\big)^2\Big], \tag{S2}$$

where the last term is the gradient penalty enforcing the 1-Lipschitz constraint on $D_\psi$, $\hat{\mathbf{x}}=\alpha\mathbf{x}+(1-\alpha)G_\theta(\mathbf{z},c)$ with $\alpha\sim\mathcal{U}(0,1)$, and $\lambda$ is the penalty coefficient.

##### Conditional RealNVP normalizing flow.

A normalizing flow (Papamakarios et al., 2021) models the data density *exactly* by pushing a simple base distribution (here a standard Gaussian) through a sequence of invertible, differentiable maps; the change-of-variables formula then gives the exact log-likelihood, so the model is trained by direct maximum likelihood rather than a bound (as in the VAE) or an implicit adversarial objective (as in the GAN). RealNVP (Dinh et al., 2017; Rezende and Mohamed, 2015) realizes this with affine coupling layers that transform one half of the input conditioned on the other, an arrangement whose Jacobian is triangular so that both the forward map and its log-determinant are cheap to evaluate. Two consequences matter here. First, exact-likelihood training over a bijective map forces the model to place probability mass over the full support of the data, including its tails, rather than collapsing onto the core. Second, this gives the flow an inductive bias toward *exploration*: it covers the reference manifold most broadly and extrapolates genuinely beyond it, which is why it most often proposes models better than the best in the initial ensemble, albeit at somewhat lower stability. Making the coupling networks' scale and translation depend on $c$ yields the conditional flow. The exact conditional log-likelihood is

$$\log p_\theta(\mathbf{x}\mid c)=\log p_{\mathbf{u}}\big(f_\theta(\mathbf{x};c)\big)+\log\left|\det\frac{\partial f_\theta(\mathbf{x};c)}{\partial\mathbf{x}}\right|, \tag{S3}$$

maximized directly over the training set, where the invertible map $f_\theta(\cdot\,;c)$ is a stack of affine coupling layers that split $\mathbf{x}$ into $(\mathbf{x}_{1:d},\mathbf{x}_{d+1:D})$ and apply $\mathbf{y}_{1:d}=\mathbf{x}_{1:d}$, $\mathbf{y}_{d+1:D}=\mathbf{x}_{d+1:D}\odot\exp(s(\mathbf{x}_{1:d},c))+t(\mathbf{x}_{1:d},c)$, with scale $s$ and translation $t$ conditioned on $c$.

##### Conditioning schemes.

Every architecture is evaluated under two conditioning (labelling) schemes that we contrast throughout. In the *discrete* scheme, $c\in\{0,1\}$ is a single per-scenario binary label separating jointly high- from low-performing ensembles at the KPI thresholds of the main text. In the *continuous* scheme, $c\in\mathbb{R}^k$ is the graded performance target itself: the vector of standardized log-KPI values, one entry per perturbation ($k$ ranging from 1 for a single mutant to 9 for the joint nine-mutant class), where each KPI is log-transformed and standardized to zero mean and unit variance using statistics fixed on the training set. The continuous scheme thus exposes the network to the full performance gradient rather than a thresholded class. The target $c$ is injected into each network through a learned embedding (discrete scheme) or by direct concatenation of the standardized KPI vector (continuous scheme). At generation time, ensembles are drawn under the high-performance regime: the positive label ($c=1$) in the discrete scheme and, in the continuous scheme, KPI target vectors taken from the discrete positive class, i.e. with every entry at or below its ORACLE class threshold, mapped through the same log-standardization, so that both schemes aim at the same performance class and the comparison between them is fair. A more demanding target, the $5^{\mathrm{th}}$ percentile of every KPI, is evaluated for Scenario III in Section S7. The generated ensembles are then passed to the stability-based filtering and KPI evaluation procedures of Section S3.

#### S3. Dynamic simulation protocol

Every generated parameter set is scored with the same pipeline used to construct the baseline ORACLE ensemble, so that all six generator variants and ORACLE are evaluated identically.

##### Stability-based screening.

Before any phenotype is evaluated, each parameter set passes the two-stage screen introduced in Section S1.1. *Dynamic stability*: the Jacobian of the ODE system is evaluated at the reference steady state, and only sets whose Jacobian eigenvalues all have strictly negative real parts are retained. *Physiological timescale*: the characteristic relaxation time, estimated from the dominant (least-negative) eigenvalue, must be at least three times faster than the doubling time $t_d=\ln 2/\mu$ ($\mu=0.2\,\mathrm{h}^{-1}$, $t_d\approx 3.47\,\mathrm{h}$ for *E. coli*), which removes models that are mathematically stable but unrealistically slow. The fraction of generated models that survive this screen is reported as the *stability* column of every leaderboard.

##### Perturbation and integration.

Each surviving model is initialized at the reference steady-state metabolite concentrations. The scenario's enzyme perturbation(s) are then imposed as multiplicative changes to the corresponding abundance parameters (Section S1.2), and the perturbed system is integrated to its post-perturbation state under quasi-steady-state conditions using the ODE solver implemented in `skimpy` (Weilandt et al., 2023). Because each single-perturbation experiment is run independently, the same screened ensemble is reused across all rungs of the difficulty ladder.

##### Phenotype scoring.

Fluxes are read out from the post-perturbation state and compared against the same experimental references (Ishii et al., 2007) used to build the baseline ensemble, through the composite ORACLE-normalized KPI ($\zeta$-score) defined in the main text. Applying an identical simulation-and-scoring procedure to ORACLE and to all three architectures under both conditioning schemes is what makes the cross-method comparison fair.

#### S4. Statistical fidelity and diversity metrics

This section gives the full definitions of the statistical fidelity and diversity metrics summarized in the main text: manifold precision and recall (Kynkäänniemi et al., 2019) and the normalized nearest-neighbour extrapolation distance. The closely related density/coverage (Naeem et al., 2020) and $\alpha$-precision/$\beta$-recall (Alaa et al., 2022) are natural alternatives.

**Feature space.**

Each raw 259-dimensional $K_m$ vector $\mathbf{k}$ is mapped to

$$\mathbf{x} \;=\; \mathrm{PCA}_L\left(\frac{\log \mathbf{k} - \boldsymbol{\mu}}{\boldsymbol{\sigma}}\right), \tag{S4}$$

where $\boldsymbol{\mu}$ and $\boldsymbol{\sigma}$ are the per-parameter mean and standard deviation of $\log K_m$ over the ORACLE reference set, and $\mathrm{PCA}_L$ projects onto the leading $L = \min(50, d, N-1)$ principal components fit on that reference. Working in this reduced space avoids the distance concentration that makes nearest-neighbour statistics unreliable in the full 259-dimensional space; all distances below are Euclidean in it. Let $R = \{\mathbf{r}_1, \ldots, \mathbf{r}_N\}$ denote the *reference* set (the ORACLE high-performing parameter sets for the scenario) and $F = \{\mathbf{f}_1, \ldots, \mathbf{f}_M\}$ the *generated* set (the surviving synthetic parameter sets).

**Precision and recall.**

For a point $\mathbf{x}$ in a set $X$, let $\mathrm{NND}_k(\mathbf{x}, X)$ be the distance to its $k$-th nearest neighbour in $X \setminus \{\mathbf{x}\}$ ($k = 5$). The estimated support of $X$ is the union of balls $\mathcal{M}(X) = \bigcup_{\mathbf{x}_i \in X} B\big(\mathbf{x}_i,\ \mathrm{NND}_k(\mathbf{x}_i, X)\big)$, and a point $\mathbf{y}$ lies in $\mathcal{M}(X)$ iff $\exists\, \mathbf{x}_i \in X :\ \|\mathbf{y} - \mathbf{x}_i\| \le \mathrm{NND}_k(\mathbf{x}_i, X)$. Precision is then the fraction of generated points inside the reference manifold (fidelity) and recall the fraction of reference points inside the generated manifold (coverage/diversity):

$$\text{precision} = \frac{1}{M} \sum_{j=1}^{M} \mathbf{1}[\ \exists\, i :\ \|\mathbf{f}_j - \mathbf{r}_i\| \le \mathrm{NND}_k(\mathbf{r}_i, R)\ ], \tag{S5}$$

$$\text{recall} = \frac{1}{N} \sum_{i=1}^{N} \mathbf{1}[\ \exists\, j :\ \|\mathbf{r}_i - \mathbf{f}_j\| \le \mathrm{NND}_k(\mathbf{f}_j, F)\ ]. \tag{S6}$$

Precision uses the neighbourhood radii of the reference points $\mathrm{NND}_k(\mathbf{r}_i, R)$, whereas recall uses those of the generated points $\mathrm{NND}_k(\mathbf{f}_j, F)$.

**Extrapolation distance.**

Precision and recall record only whether a sample falls inside or outside the reference manifold; to capture the *magnitude* of the departure we also report a normalized nearest-neighbour distance. For a generated model $\mathbf{f}_j$ we take the mean distance to its $k$ nearest reference points, $d_k(\mathbf{f}_j, R) = \frac{1}{k} \sum_{\mathbf{r} \in \mathcal{N}_k(\mathbf{f}_j, R)} \|\mathbf{f}_j - \mathbf{r}\|$, and normalize by the reference set's own internal scale $s_R = \mathrm{median}_{\mathbf{r} \in R}\ d_k(\mathbf{r}, R \setminus \{\mathbf{r}\})$, giving the per-model ratio and its ensemble mean

$$\rho_j = \frac{d_k(\mathbf{f}_j, R)}{s_R}, \qquad \bar{\rho}(F, R) = \frac{1}{M} \sum_{j=1}^{M} \rho_j. \tag{S7}$$

The ratio reads directly: $\rho \approx 1$ denotes interpolation within the reference (a model as far from it as its members are from one another), $\rho \gg 1$ genuine extrapolation beyond the sampled manifold, and $\rho \approx 0$ near-duplication of reference models (memorization). We report $\bar{\rho}$ against two references: the full ORACLE ensemble, quantifying overall novelty, and the scenario-specific high-performing subset $R$, quantifying extrapolation beyond the known good region and hence the capacity to propose parameter sets superior to those already sampled. All statistical metrics are reported as the mean ± standard deviation across random seeds.

#### S5. Scenario I (PGI) full leaderboard

Table S4 reports the full Scenario I leaderboard summarized in the main text.

#### S6. Additional figures

Figure S1 details the per-parameter $\log(K_m)$ shifts for Scenario I; Figure S2 shows the per-perturbation KPI densities for Scenario II; Figure S3 shows the per-mutant densities for the stringent nine-perturbation class; and Figure S4 shows the Scenario I extrapolation distances.

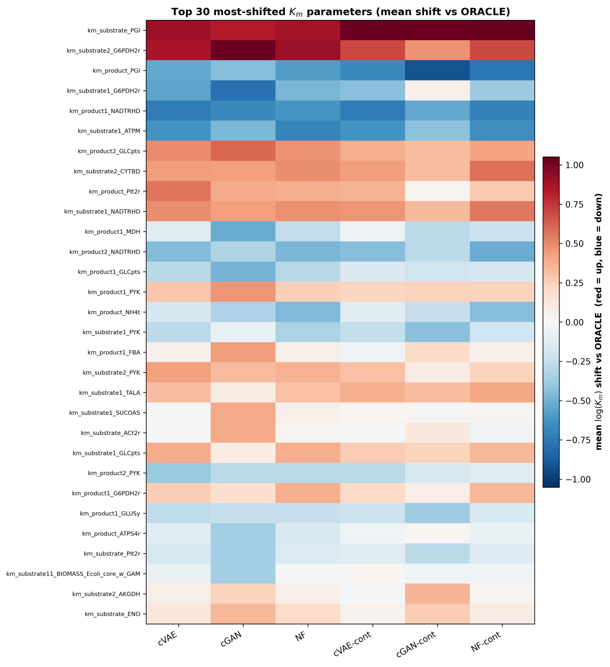


**Figure S1 Scenario I (PGI): mean $\log(K_m)$ shift relative to ORACLE** (red up, blue down), top-30 parameters. The sign pattern is shared across variants (common manifold); only the discrete cGAN flips a few (realigns under cGAN-cont).

#### S7. Scenario III: targeted generation beyond the positive class

The main-text comparison conditions the continuous variants on targets drawn from the positive class, so that they aim at the same class as the discrete label. Because the continuous target is graded, it can also be set below that class. Here the three continuous variants are conditioned on a single target vector holding the $5^{\mathrm{th}}$ percentile of each of the nine ORACLE KPIs, mapped through the same log-standardization; all other settings, and the discrete variants, are

**Table S4** Scenario I (PGI) leaderboard; mean ± s.d. over seeds ($n = 4$; $n = 3$ for cGAN-cont). In-thr.: fraction below ORACLE p10; Best-1%: fraction in lowest 1% of ORACLE KPI; Beat-best: fraction beating the best ORACLE model; Stable: stability pass rate; Rec.: manifold coverage (Kynkäänniemi et al., 2019); $\bar{\rho}_{\mathrm{full}}$: mean NN distance to ORACLE (≈1 interpolate, >1 extrapolate). Best per column (excl. ORACLE) in bold.

| Ensemble | Mean KPI | In-thr. (%) | Best-1% (%) | Beat-best (%) | Stable (%) | Rec. | $\bar{\rho}_{\mathrm{full}}$ |
|---|---|---|---|---|---|---|---|
| ORACLE | 12.05 | 10.0 | 1.0 | 0.00 | 62.8 | – | 1.00 |
| cVAE | **7.45** ± 0.17 | **87.5** ± 5.7 | **11.8** ± 3.1 | 0.00 ± 0.00 | **100.0** ± 0.0 | 0.00 | 0.77 |
| cGAN | 9.67 ± 0.27 | 38.3 ± 3.0 | 7.1 ± 0.8 | 0.08 ± 0.06 | 78.9 ± 3.4 | 0.04 | 0.99 |
| NF | 9.73 ± 0.10 | 36.8 ± 1.2 | 6.2 ± 0.8 | **0.14** ± 0.05 | 73.7 ± 0.8 | 0.37 | **1.08** |
| cVAE-cont | 7.53 ± 0.15 | **88.0** ± 7.0 | 4.1 ± 1.8 | 0.00 ± 0.00 | **100.0** ± 0.0 | 0.00 | 0.77 |
| cGAN-cont | 9.43 ± 0.53 | 32.8 ± 9.7 | 3.3 ± 1.5 | 0.00 ± 0.00 | 79.4 ± 1.8 | 0.03 | 0.98 |
| NF-cont | 9.27 ± 0.10 | 37.6 ± 1.5 | 4.7 ± 0.4 | 0.07 ± 0.08 | 74.7 ± 0.4 | 0.37 | **1.08** |

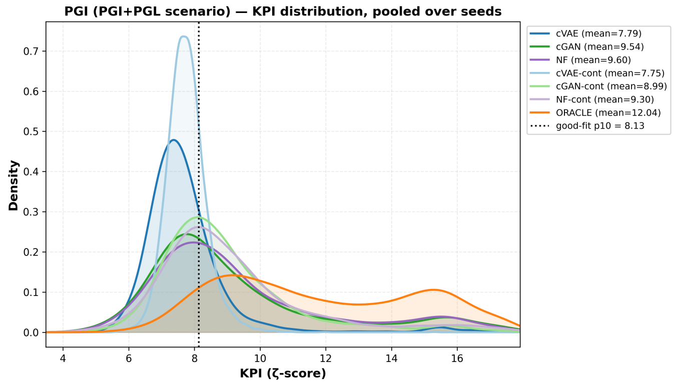


(a)

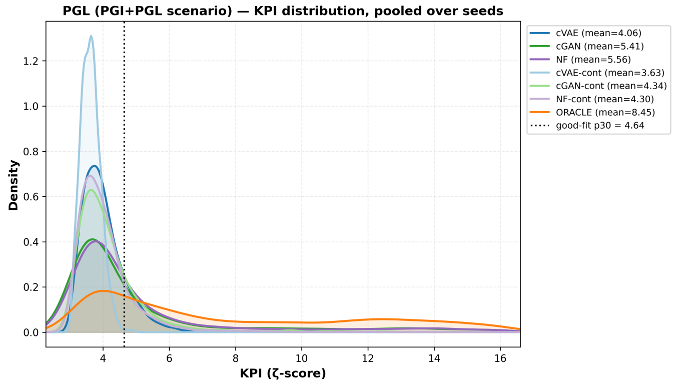


(b)

**Figure S2 Scenario II (PGI+PGL): per-perturbation KPI ($\zeta$-score) densities**, pooled over seeds (dotted: good-fit partition, PGI p10 / PGL p30; panels share one legend). **(a)** PGI and **(b)** PGL. All variants shift left of ORACLE, most sharply the cVAE.

unchanged. The more demanding target raises the joint-class yield 2.9–3.6× for every architecture (Table S5, Fig. S5), while recall and the extrapolation distance $\bar{\rho}_{\mathrm{full}}$ remain essentially unchanged: the ensembles concentrate further within the known high-performing support rather than moving into new regions of parameter space.

D. J. Rezende and S. Mohamed. Variational inference with normalizing flows. In *International Conference on Machine*

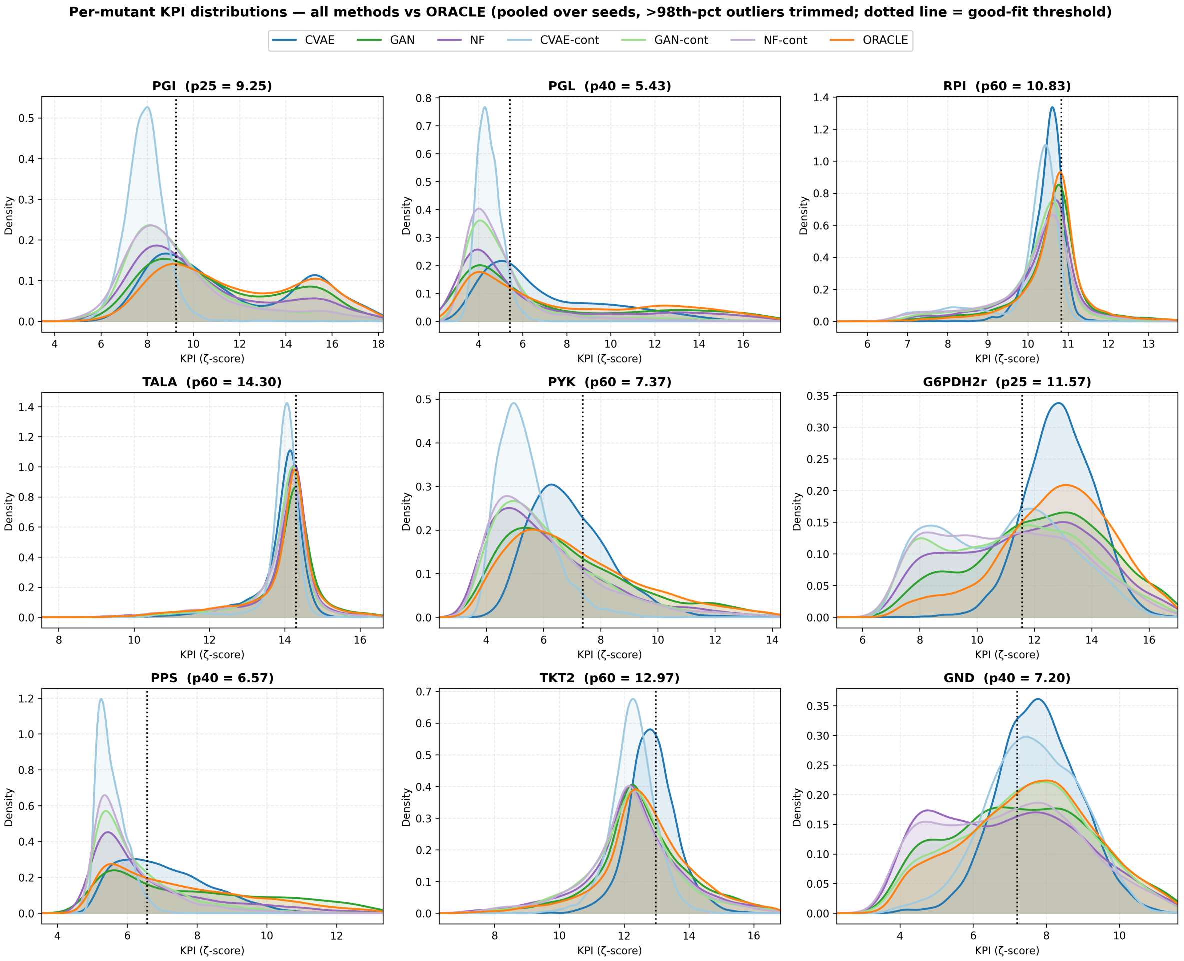


**Figure S3 Scenario III (stringent 9-mutant joint class): per-mutant KPI ($\zeta$-score) densities**, pooled over seeds (dotted: good-fit threshold); continuous variants conditioned on positive-class targets. Continuous variants pull mass below threshold on nearly every mutant at once (the exception is cVAE-cont on GND, $36\%$ below threshold versus $40\%$ for ORACLE); discrete variants clear the easy mutants but miss the hard ones, which is what makes the joint all-nine class so much harder for them.

**Table S5** Scenario III continuous variants under the positive-class target (main text) and the $5^{\text{th}}$-percentile target; mean $\pm$ s.d. over seeds ($n = 3$). Joint %: fraction satisfying all nine thresholds; Enrich.: over the ORACLE base rate $(0.13\%)$; $\geq 7/9$: fraction satisfying at least seven of the nine thresholds; Mean KPI: averaged over the nine perturbations (non-physical KPIs $> 10^3$ excluded); Stable: stability pass rate; Rec.: manifold coverage; $\bar{\rho}_{\text{full}}$: mean NN distance to ORACLE; Beat-best: perturbations (of nine) with at least one model beating the best ORACLE model. For the $5^{\text{th}}$-percentile target, cVAE-cont and NF-cont use seeds $45$, $66$, $67$ and cGAN-cont seeds $65$–$67$; all positive-class runs use seeds $65$–$67$.

| Variant | Target | Joint (%) | Enrich. $\times$ | $\geq 7/9$ (%) | Mean KPI | Stable (%) | Rec. | $\bar{\rho}_{\text{full}}$ | Beat-best |
|---|---|---|---|---|---|---|---|---|---|
| cVAE-cont | positive class | $15.0 \pm 4.4$ | $117 \pm 34$ | $85.2 \pm 2.6$ | 8.66 | $99.9 \pm 0.1$ | 0.00 | 0.77 | $0.0 \pm 0.0$ |
| | $5^{\text{th}}$ pct. | $53.6 \pm 23.3$ | $416 \pm 181$ | $99.5 \pm 0.5$ | 7.91 | $99.9 \pm 0.0$ | 0.00 | 0.76 | $0.0 \pm 0.0$ |
| cGAN-cont | positive class | $4.2 \pm 0.9$ | $33 \pm 7$ | $40.3 \pm 3.9$ | 9.34 | $75.7 \pm 7.1$ | 0.03 | 1.00 | $2.3 \pm 1.2$ |
| | $5^{\text{th}}$ pct. | $12.3 \pm 5.1$ | $95 \pm 40$ | $68.6 \pm 5.3$ | 8.70 | $69.9 \pm 10.6$ | 0.01 | 0.99 | $1.7 \pm 0.6$ |
| NF-cont | positive class | $5.6 \pm 1.0$ | $43 \pm 8$ | $45.7 \pm 3.0$ | 9.32 | $70.3 \pm 1.6$ | 0.33 | 1.09 | $2.3 \pm 0.6$ |
| | $5^{\text{th}}$ pct. | $16.5 \pm 2.4$ | $128 \pm 19$ | $74.9 \pm 1.8$ | 8.77 | $66.7 \pm 3.8$ | 0.36 | 1.11 | $3.3 \pm 1.5$ |

*Learning (ICML)*, pages 1530–1538, 2015.

K. Sohn, X. Yan, and H. Lee. Learning structured output representation using deep conditional generative models. In *Advances in Neural Information Processing Systems (NeurIPS)*, volume 28, 2015.

D. R. Weilandt, P. Salvy, M. Masid, G. Fengos, R. Denhardt-Eriksson, Z. Hosseini, and V. Hatzimanikatis. Symbolic kinetic models in Python (SKiMpy): intuitive modeling of large-scale biological kinetic models. *Bioinformatics*, 39(1):btac787, 2023.

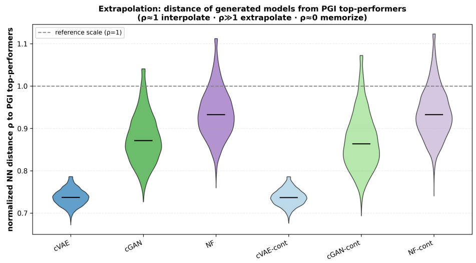


**Figure S4 Scenario I (PGI): nearest-neighbour distance $\rho$ to the PGI top-performers**, pooled over seeds ($\rho \approx 1$ interpolation, $> 1$ extrapolation): the cVAE contracts, the flows extrapolate, and the cGAN variants lie between.

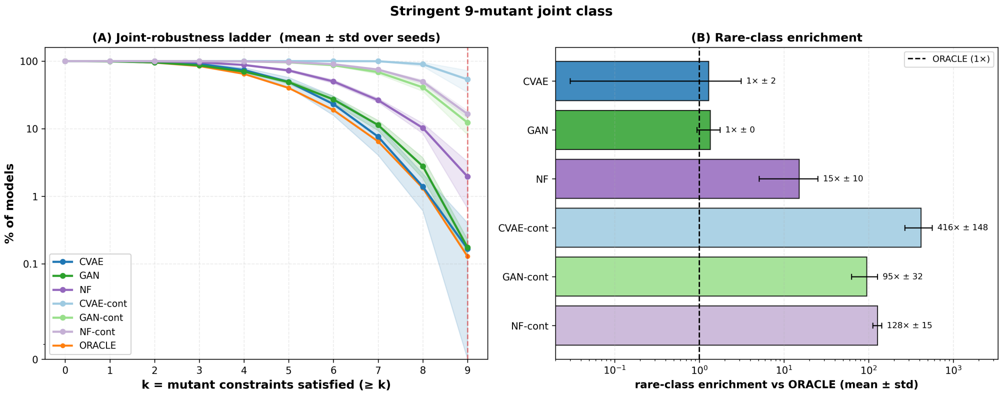


**Figure S5 Scenario III with the $5^{\mathrm{th}}$-percentile continuous target**, pooled over seeds. Joint-robustness ladder (% of models satisfying $\geq k$ of $9$ thresholds) and rare-class enrichment vs. ORACLE; discrete variants as in main-text Fig. 4. Compared with the positive-class target, every continuous variant moves further up the ladder (cVAE-cont $416\times$, NF-cont $128\times$, cGAN-cont $95\times$).
